# Targeting current species ranges and carbon stocks fails to conserve biodiversity in a changing climate: opportunities to support climate adaptation under 30×30

**DOI:** 10.1101/2021.08.31.458416

**Authors:** Lindsay M. Dreiss, L. Mae Lacey, Theodore C. Weber, Aimee Delach, Talia E. Niederman, Jacob W. Malcom

**Affiliations:** Center for Conservation Innovation, Defenders of Wildlife, Washington, DC 20036, USA; Department of Landscape Conservation, Defenders of Wildlife, Washington, DC 20036, USA

**Keywords:** Climate refugia, Climate corridors, Protected areas, Biodiversity conservation, Carbon mitigation

## Abstract

Protecting areas for climate adaptation will be essential to ensuring greater opportunity for species conservation well into the future. However, many proposals for protected areas expansion focus on our understanding of current spatial patterns, which may be ineffective surrogates for future needs. A science-driven call to address the biodiversity and climate crises by conserving at least 30% of lands and waters by 2030, 30×30, presents new opportunities to inform the siting of new protections globally and in the U.S. Here we identify climate refugia and corridors based on a weighted combination of currently available models; compare them to current biodiversity hotspots and carbon-rich areas to understand how 30×30 protections siting may be biased by data omission; and compare identified refugia and corridors to the Protected Areas Database to assess current levels of protection. Available data indicate that 20.5% and 27.5% of identified climate adaptation areas (refugia and/or corridor) coincides with current imperiled species hotspots and carbon-rich areas, respectively. With only 12.5% of climate refugia and corridors protected, a continued focus on current spatial patterns in species and carbon richness will not inherently conserve places critical for climate adaptation. However, there is ample opportunity for establishing future-minded protections: 52% of the contiguous U.S. falls into the top quartile of values for at least one class of climate refugia. Nearly 27% is already part of the protected areas network but managed for multiple uses that may limit their ability to contribute to the goals of 30×30. Additionally, nearly two-thirds of nationally identified refugia coincide with ecoregion-specific refugia suggesting representation of nearly all ecoregions in national efforts focused on conserving climate refugia. Based on these results, we recommend that land planners and managers make more explicit policy priorities and strategic decisions for future-minded protections and climate adaptation.

## INTRODUCTION

The spatial heterogeneity of shifting climatic conditions presents challenges and opportunities for large-scale biodiversity conservation, as impacts to habitat and species can vary significantly across the landscape (Baldwin et al. 2018). In North America, nearly half of species are already undergoing local extinctions (Wiens 2016), which are partially due to spatially variable changes in temperature and precipitation (Román-Palacios and Wiens 2020). In the contiguous U.S. (CONUS), the average annual temperature has risen 1.2-1.8 °C since the beginning of the 20th century (Vose et al. 2017) and precipitation patterns have shifted with large reductions in the Southeast and West (Fei et al. 2017, Wuebbles et al. 2017). As the effects of climate change accelerate, local biodiversity will either need to find locations that serve as refugia from extreme or rapid climatic changes or shift their ranges to better-suited habitat (Neilson et al. 2005, Keppel and Wardell-Johnson 2012, Franks and Hoffman 2012, Román-Palacios and Wiens 2020). Identifying and conserving important refugia habitats and dispersal routes will be one critical step in jointly addressing the biodiversity and climate crises (Pörtner et al. 2021). While expansion of the U.S. protected areas network has been identified as an important solution to lowering extinction risk and overall ecosystem degradation (Stolton et al. 2015, Gray et al. 2016, Dinerstein et al. 2017, 2019), efforts generally focus on current species distributions and may not effectively reflect future needs (Elsen et al. 2020, Maxwell et al. 2020).

Calls to address the joint biodiversity and climate crises by protecting at least 30% of Earth by 2030, known as “30×30” (Dinerstein et al. 2019), have been endorsed by government and conservation leaders at global (United Nations 2020), national (Biden 2021, U.S. DOI et al. 2021), and state levels (e.g., Newsom 2020). While the specifics have yet to be established (Büscher et al. 2016, Rights and Resources Initiative 2020, Simmons et al. 2021), efforts would hypothetically conserve areas needed to sustain essential ecological services and reverse extinction trends (Locke 2013, Dinerstein et al. 2017). Translating these commitments into national policy may prove challenging since the current distribution of protected areas excludes many areas important for biodiversity conservation (Scott et al. 2001, Jenkins et al. 2015, Venter et al. 2018) and carbon storage (Buotte et al. 2019, Melillo et al. 2015). While not all areas storing large amounts of carbon are also biodiversity hotspots, there are many instances of overlap; for instance, carbon-rich forests can provide important habitat and climate change buffers (Berkessy and Wintle 2008, Strassburg et al. 2010). However, it is unclear how well the current network and 30×30 goals can ensure the conservation of climate-resilient habitat in the coming decades as climate change continues to accelerate.

Climate-resilient habitat can largely be delineated into refugia and corridors. Refugia are areas where species, natural communities, or ecosystems can persist within a larger area that has been rendered inhospitable. Such areas are relatively buffered from changes in regional environmental conditions, and allow organisms to persist as long as local conditions remain tolerable (Keppel and Wardell-Johnson, 2012; Morelli et al., 2016). Climate-change refugia may only be temporary for a given species (Morelli et al., 2020), but can delay ecosystem transitions for decades or longer (Krawchuk et al., 2020). They can serve as a “slow lane,” protecting native species and ecosystems from negative effects of climate change in the short term, providing longer-term havens for overall biodiversity and ecosystem function, and reducing the risk of extinction (Morelli et al., 2020). Although refugia can be examined over a continuum of spatial scales, they are usually classified as either macrorefugia or microrefugia (Ashcroft, 2010). Macrorefugia are identified at coarse scales, using global climate data or models, and are large enough to maintain viable animal or plant populations. Microrefugia are identified at local scales. For example, steep canyons and north-facing slopes are sheltered from solar radiation and heat accumulation (Stralberg et al., 2020a) and wet areas can remain moist during droughts (Morelli et al., 2016; Stralberg et al., 2020a).

At broad scales, refugia can be identified by various approaches, such as topodiversity, climate exposure, and climate tracking (Michalak et al. 2020). Topodiversity models are based on physical habitat data and highlight regions with varied land cover, climate, soil, and topographic conditions (Ackerly et al. 2010, Groves et al. 2012, Carroll et al. 2018). Topographically varying areas can contain features like deep valley bottoms or shaded slopes that serve as microrefugia (Ashcroft 2010). Climatic exposure models are based on projected climatic changes and represent the degree of climate change likely to be experienced by a species or locale (Saxon 2011, Groves et al. 2012). Lastly, climate tracking models are based on representative climate models and measure the proximity and accessibility of future suitable climatic conditions (Hamann et al. 2015, Michalak et al. 2018).

However, to survive in the face of ongoing and worsening climate change impacts, some species may need to relocate (Román-Palacios and Wiens 2020). Climate corridors can facilitate long-distance movement to more suitable habitat if rising temperatures and other changing conditions exceed an organism’s tolerance (Stralberg et al., 2020b). Depending on model inputs, corridors may emphasize movement toward cooler latitudes and topographies, along rivers and streams, and/or through areas providing better habitat and less stress from disturbances (McGuire et al. 2016, Stralberg 2020b, Carroll et al. 2018, Littlefield et al. 2017). For example, Parks et al. (2020) found that human land modifications decrease climate connectivity.

Given the urgency of the biodiversity and climate crises, there is a pressing need to include potential climate refugia and corridors in the conservation planning process. However, some challenges exist. First, a growing body of available spatial data for identifying areas important for climate adaptation means that planners must reconcile a diversity of data that represent different mechanisms and priorities (Michalak et al. 2020, Carroll and Ray 2021). Second, most conservation prioritization frameworks focus on the current state of species and environments (Cushman et al. 2009, Lookingbill et al. 2010, Dickson et al. 2013, Belote et al. 2016, McClure et al. 2016) which may result in critical omissions in protected areas siting for longer-term persistence of some target species (Monzón et al. 2011, Elsen et al. 2020). If this is the case, consideration of future conditions may complement efforts to preserve current biodiversity and ecosystem service hotspots, thereby reducing the threat of mass extinctions and accompanying biosphere degradation. Last, other omissions of important local refugia may occur if areas for climate resilience are identified and prioritized at a national scale (e.g. Kraus and Hebb 2020). While this may identify the most important nationwide climate refugia, it could favor some regions (e.g., mountain ranges) and omit others entirely. Evaluating at smaller scales (e.g., stratifying by ecoregion) could include more locally important refugia. Taking additional steps to identify refugia at multiple scales may help increase ecosystem representation and protections for the unique species assemblages and services they harbor.

Proper identification, protection, and management of climate-informed refugia and corridors are essential to ensuring greater opportunity for species conservation via migration and adaptation. While previous research and policy discussion surrounding the extent and distribution of protected land and water has identified areas important to conserving the current state of biodiversity and natural carbon storage (Scott et al. 2001, Myers et al. 2000, Gray et al. 2016, Buotte et al. 2020), less attention has been paid to areas important for wildlife and plant climate adaptation. To help close this knowledge gap, we:

1. identify areas in the contiguous U.S. critical to climate adaptation based on coincidence and complementarity among refugia (national and ecoregion-specific) and corridors models;
2. compare the spatial distribution of identified climate refugia and corridors with current biodiverse and carbon-rich areas; and
3. quantify the extent to which climate refugia and corridors are considered protected.

Given the opportunity to focus land conservation using a science-driven, ecologically-meaningful (rather than opportunistic) approach, step #2 guides our understanding of how protections siting under the 30×30 framework may be biased by data omission, and step #3 helps to assess the current level of protection for identified climate refugia and distinguish where stronger management might be needed. Our research contributes to a growing literature demonstrating the importance of incorporating climate-informed data in place-based land protection policy and practices and helping to identify specific areas for conservation. While these analyses are not meant to serve as a map of priority lands for conservation, they help guide discussion on operationalizing 30×30 for strategic, future-minded conservation decisions.

## METHODS

We focus on spatial datasets based on climate models or topography to identify areas that could serve as important refugia or migration routes for the contiguous U.S. (CONUS; Table 1). All datasets using climate models are informed by an ensemble of three to seven General Circulation Models (GCMs) for emission scenario Representative Concentration Pathway (RCP) 4.5 and projected for the time period 2071-2100 based on the Coupled Model Intercomparison Project phase 5 (CMIP5) models. All datasets have been resampled and aligned at 1km resolution. We combined datasets for refugia (n = 8) and corridors (n = 2) separately, accounting for differences in underlying mechanisms in modeling method and landscape conservation principles.

**Table 1.**
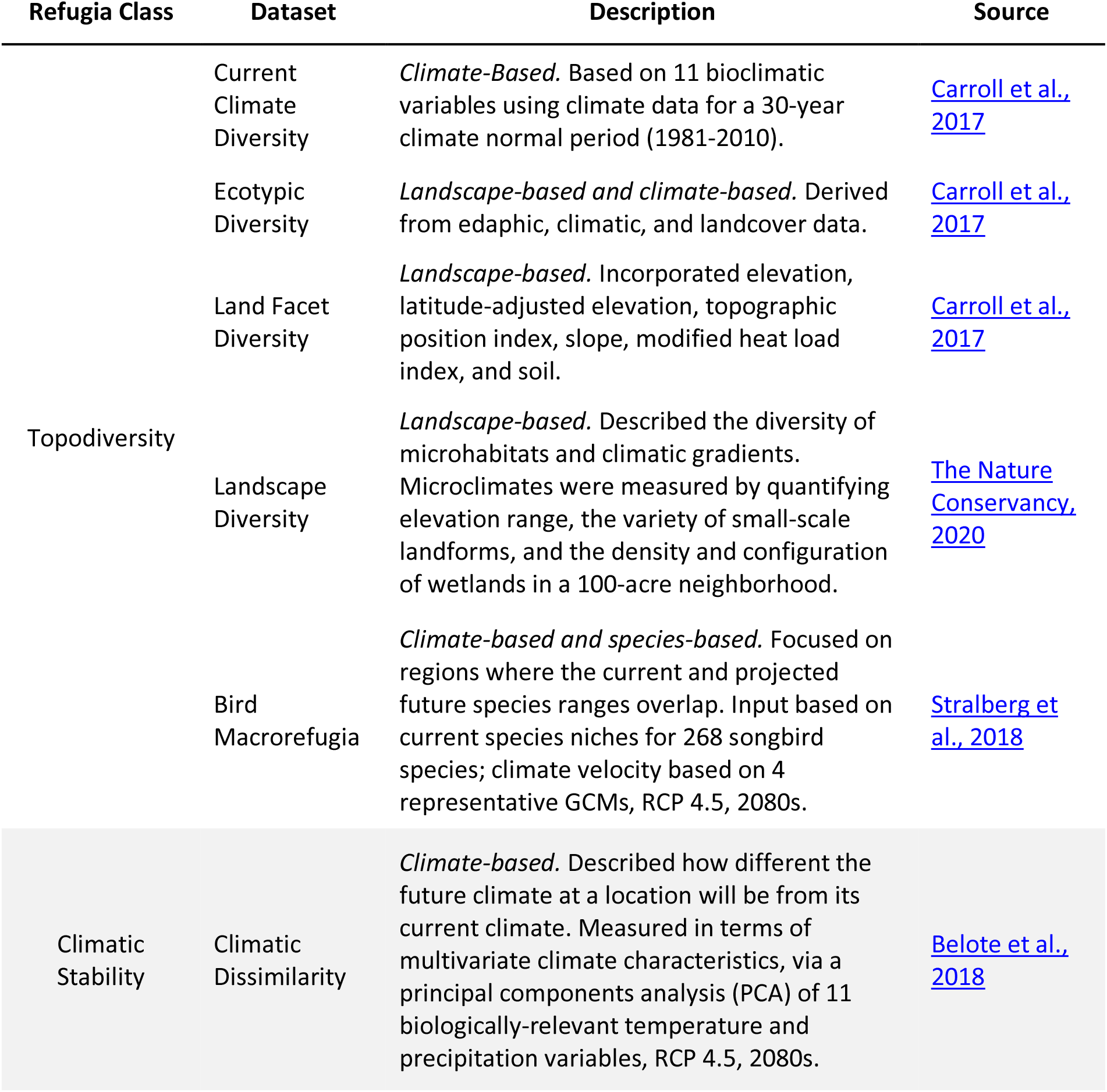
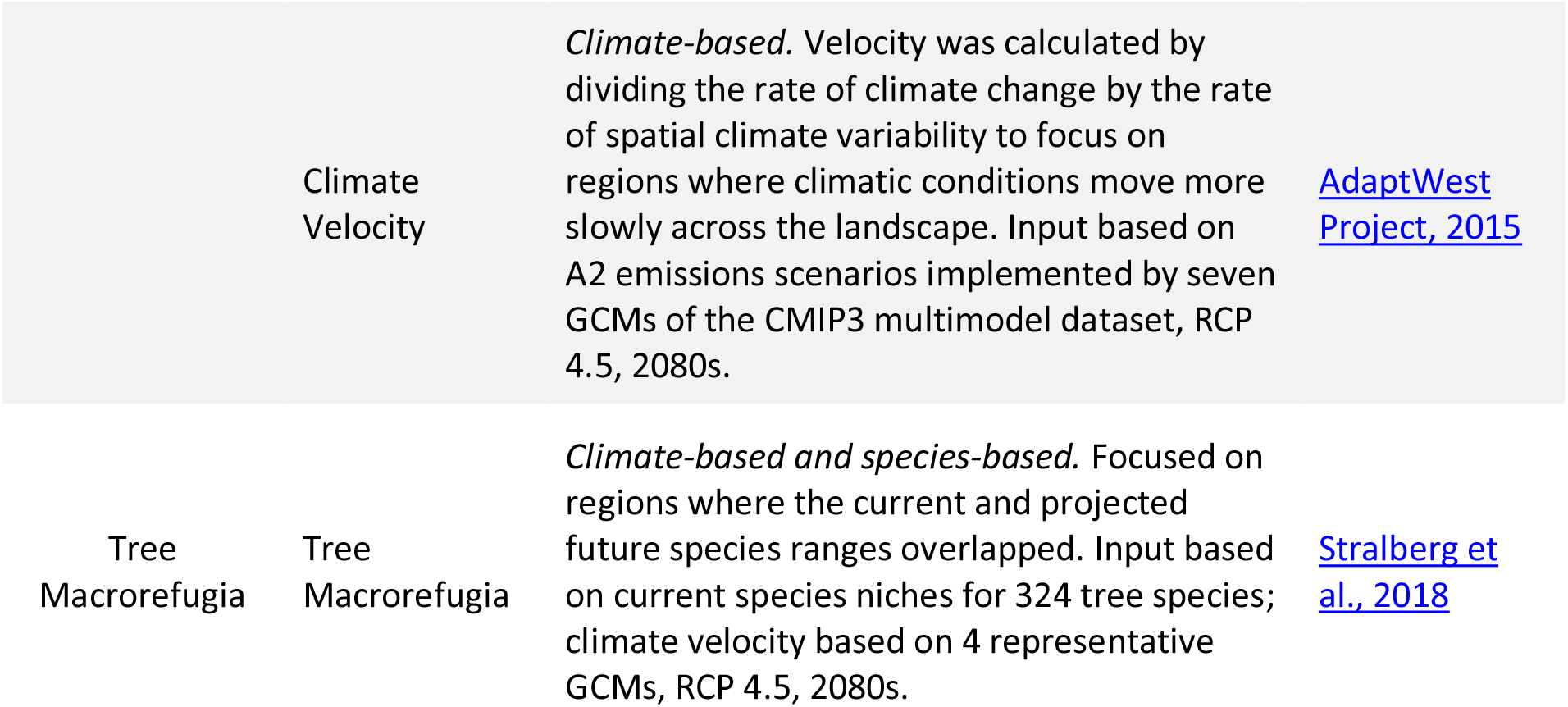
Description of refugia datasets. All datasets are currently based on the Coupled Model Intercomparison Project phase 5 (CMIP5) models. Classes are based on results from a principal components analysis where component 1 (topodiversity) explained 33.8%, component 2 (climate stability) explained 15.9% and component 3 (tree macrorefugia) explained near 13.8% of variation. See SI for additional details.

### Climate refugia

We initially analyzed relationships between datasets through a principal components analysis where each component helps define a refugia class. The principal components spatial analyst tool in ArcPro transforms the data in the input raster bands to compress data and eliminate redundancy. The result is a multiband raster and text output containing a covariance matrix, correlation matrix, eigenvalues and eigenvectors. The number of principal components selected was based on scree plot and broken stick model. All datasets were normalized to a scale of 0 to 1 by calculating z-scores for all eight refugia datasets representing climate-, landscape-, and species-based metrics (Table 1), which were then used as inputs for the principal components analysis.

As with principal components, datasets were assigned to a class based on the sign and size of the eigenvector. However, to avoid a tradeoff in refugia identification within a single class, all datasets within the class were required to load together and in the same direction on a principal component. In addition to presenting three separate classes that represented topodiversity, climatic stability, and tree macrorefugia, we weighted all eight normalized datasets such that each dataset contributed equally to its class and each class contributed equally to the final pixel value. Unlike existing studies in this area (e.g., Michalak et al. 2020), this approach groups datasets together by relationship and reduces the chances of over-emphasizing any one variable in identifying refugia. Based on the relationships between refugia datasets, the weighted combination was calculated as follows, where each variable in the equation represents the normalized spatial dataset:

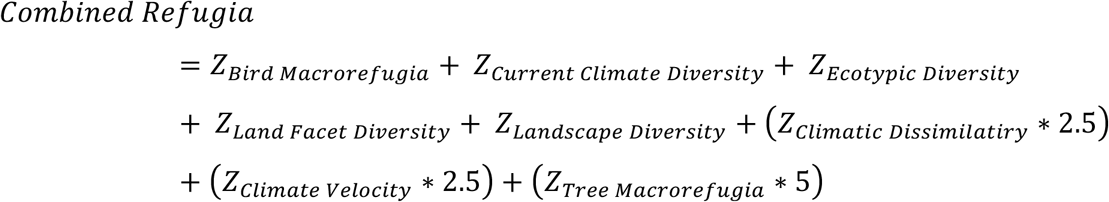

We analyzed locations in the 80^th^ percentile (i.e., the top 20% of values) of the distribution of values for the combined data and for each refugia class separately. Additionally, we quantified the degree of overlap in refugia classes.

In addition to CONUS-level analyses, we extracted refugia values for each ecoregion separately (EPA level II; EPA 2006), classifying the locations that fell into the top 20% of the distribution as areas of interest. The result was a map of ecoregion-specific refugia, ensuring equal representation of all ecoregions relative to size. Results from the national- and ecosystem-scale analyses were compared and contrasted using spatial overlays.

### Climate corridors

We extracted raw data values on connectivity and climate flow (The Nature Conservancy 2020) for areas that were identified as ‘climate-informed’ corridors by using the categorical connectivity and climate flow dataset as a mask (The Nature Conservancy 2020). The remaining values were rescaled to fall between 0 and 1. A second climate corridor dataset on current flow centrality (Carroll et al. 2018) was similarly rescaled. We then combined these two datasets and analyzed locations in the 80^th^ percentile of the distribution of combined values.

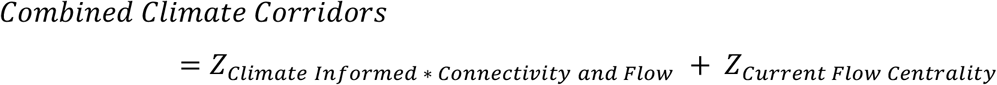

### Analyses

We used spatial overlay analysis to describe the extent to which the current protected areas network covers identified climate refugia (based on national- and ecoregion-scales) and corridors in CONUS. We quantified the extent to which identified refugia would be protected by the 30×30 framework if it were to solely focus on current areas of high imperiled species biodiversity and ecosystem carbon. Data on protected areas are from the PADUS 2.1 database (USGS 2020). We use U.S. Geological Survey’s Gap Analysis Program (GAP) codes, which are specific to the management intent to conserve biodiversity. GAP 1 and 2 areas are managed in ways typically consistent with conservation. Areas assigned a GAP 3 code are governed under multiple-use mandates that may include biodiversity priorities but may also include incompatible activities such as forestry and mining, and GAP 4 areas lack any conservation mandates or such information is unknown as of 2020. Imperiled species richness was assessed from publicly available range data (USGS GAP, International Union of Conservation of Nature - IUCN, and U.S. Fish and Wildlife Service) for species defined as ‘imperiled’ (1,923 species). These include species that are listed or under consideration for listing under the ESA, have a NatureServe G1-3 status and/or are in critically endangered, endangered or vulnerable IUCN categories. Modeled current total ecosystem carbon is based on a high-resolution map of global above- and below-ground carbon stored in biomass and soil (Soto-Navarro et al. 2020). We define hotspots for biodiversity and carbon as the top 20% of values in the raw distribution of each dataset. We selected the top 20% as a threshold to balance selectivity and breadth of coverage, but recognize that other thresholds could be chosen. We used ArcPro v2.5 (ESRI, USA) to produce maps and run analyses, with maps using the Albers Equal Area Conic projection.

## RESULTS

### Identifying refugia and corridors

Climate refugia datasets generally correlated well with others of similar methodology or concept; three resulting classes generally represent topodiversity, climatic stability, and tree macrorefugia (Tables 1 & S1). The main exception was for climate-based datasets with species information, where bird macrorefugia correlated with datasets based on topodiversity, but tree macrorefugia was the sole dataset in its class (Table S2). The three refugia classes exhibited very little overlap with one another at the national scale: while 52% of CONUS falls into at least one of the refugia classes, 7.5% falls into refugia identified by 2 or more classes (approx. 568,000 km^2^, Fig. S1). Locations in the combined refugia layer that were within the top 20% of the distribution of values represent these overlaps and are used for reporting the remainder of statistics here.

34% of CONUS is identified as a climate refugia or corridor under one or more datasets (approx. 2,652,000 km^2^, Fig. 1). Climate refugia generally follow the Appalachian, Rocky, and Cascade Mountain Ranges with additional refugia in the Ozarks and parts of California. Climate corridors are somewhat complementary to national-scale refugia, with 28.9% of their area (444,501 km^2^) overlapping identified refugia locations. Overlaps occur in the central Appalachians, Pacific Northwest, and portions of the Rockies, Sierra Nevadas and Ozarks. Corridors that do not overlap with refugia are key in connecting parts of the Great Plains and Mexico borderlands to refugia and in connecting refugia to northern locales.

**Figure 1.**
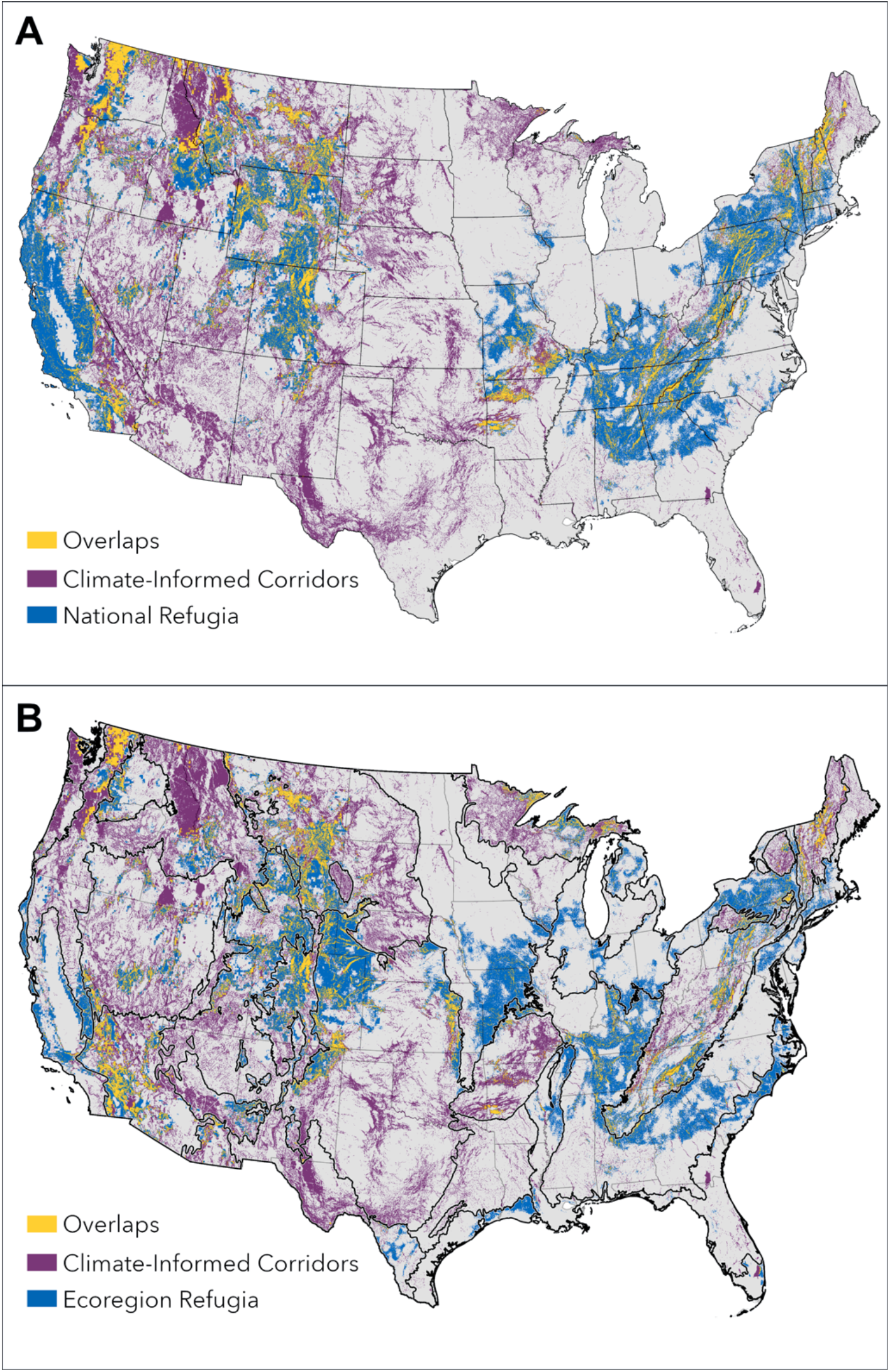
A) National-scale and B) ecoregion-specific refugia (top 20% of all three refugia classes combined) with climate-informed corridors (ecoregions are outlined in black). The full raster datasets were used to identify refugia in national analyses. Ecoregion-specific analyses employ a stratified approach, where refugia are identified for each ecoregion separately before combining them together. Ecoregions are outlined in black in map B.

Refugia identified in a stratified ecoregion approach were highly coincident with the national scale analysis, with 63% of all national refugia overlapping with ecoregion refugia (Fig. 2). Overlaps between the two cover 12% of CONUS total land area (approx. 949,000 km^2^). All refugia combined (both from national and ecoregion-specific analyses) equal 26% of the total CONUS land area (approx. 2.1 million km^2^). Locations that were emphasized in the ecoregion-specific approach include temperate and semi-arid prairies and places along the eastern coast.

**Figure 2.**
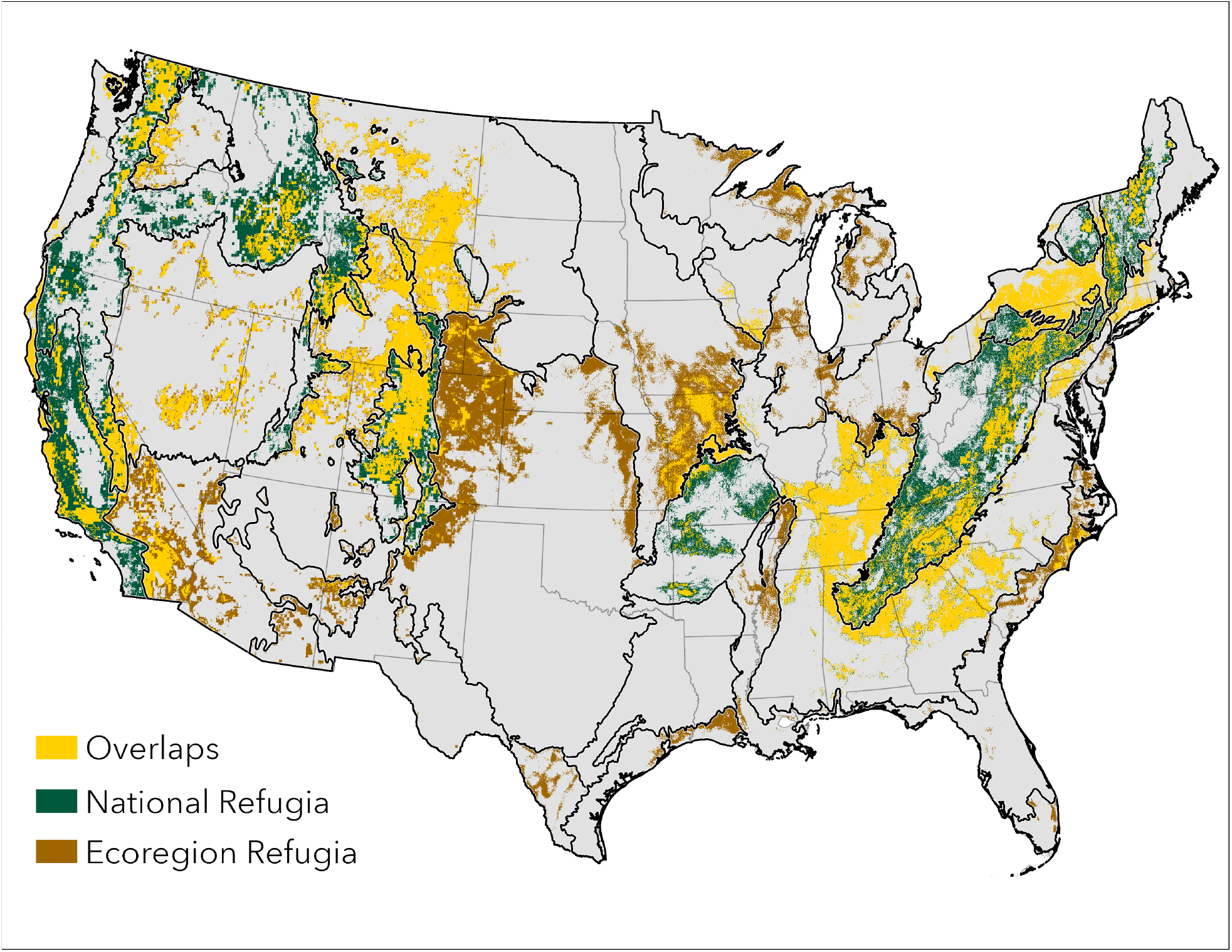
Coincidence between national-scale and ecoregion-specific refugia. The full raster datasets were used to identify refugia in national analyses. Ecoregion-specific analyses employ a stratified approach, where refugia are identified for each ecoregion separately before combining them together. Ecoregions are outlined in black.

### Comparison to 30×30 objectives: biodiversity and carbon

Refugia and corridors are generally complementary on the landscape to areas of current high biodiversity and carbon storage values (Fig. 3a&b). There is some overlap between current biodiversity hotspots (i.e., top quartile of imperiled species richness values) and identified national-scale refugia (36.8%) and corridors (9.3%; Table 2). Overlaps are generally concentrated in western California and Appalachia/Ozarks regions. Overlap between carbon-rich areas is greater in extent overall (refugia overlap = 32.5% and corridor overlap = 27.2%) and similar in spatial pattern with greater overlap in northern areas: northern Appalachians, Crown of the Continent and Pacific Northwest. When combining the two objectives (biodiversity and/or carbon), 45.0% (approx. 1,000,000 km^2^) of the land area representing at least one of these objectives is also identified as part of a climate refuge or corridor.

**Table 2.** Overlays of national-level datasets representing protected areas, carbon stores, biodiversity, climate refugia, and climate corridors. Values represent the percent of each top line item (column) that falls within each row. Values in parentheses are the percent of total CONUS area represented by the overlay.

| *% of top line items*<br>*that fall into each*<br>*of the following:* | **GAP 1 & 2** | **GAP 3** | **Top 20%**<br>**Carbon** | **Top 20%**<br>**Biodiversity** | **Top 20%**<br>**Refugia** | **Top 20%**<br>**Climate-**<br>**Informed**<br>**Corridors** |
| --- | --- | --- | --- | --- | --- | --- |
| **GAP 1 & 2** | 100 (7.5) | 0.0 (0.0) | 12.7 (2.4) | 3.7 (0.7) | 13.3 (2.6) | 13.8 (2.7) |
| **GAP 3** | 0.0 (0.0) | 100 (16.6) | 20.2 (3.9) | 5.1 (1.0) | 25.0 (4.8) | 30.4 (6.0) |
| **Top 20% Carbon** | 32.8 (2.4) | 23.3 (3.9) | 100 (20.0) | 28.8 (5.6) | 32.5 (6.2) | 27.2 (5.4) |
| **Top 20%**<br>**Biodiversity** | 12.3 (0.7) | 8.7 (1.0) | 32.0 (5.6) | 100 (20.0) | 30.8 (5.9) | 11.2 (1.8) |
| **Top 20% Refugia** | 34.2 (2.6) | 29.0 (4.8) | 32.8 (6.2) | 36.8 (5.9) | 100 (20.0) | 28.7 (5.7) |
| **Top 20% Climate-**<br>**Informed**<br>**Corridors** | 36.5 (2.7) | 36.3 (6.0) | 25.8 (5.4) | 9.3 (1.8) | 29.6 (5.7) | 100 (20.0) |

**Figure 3.**
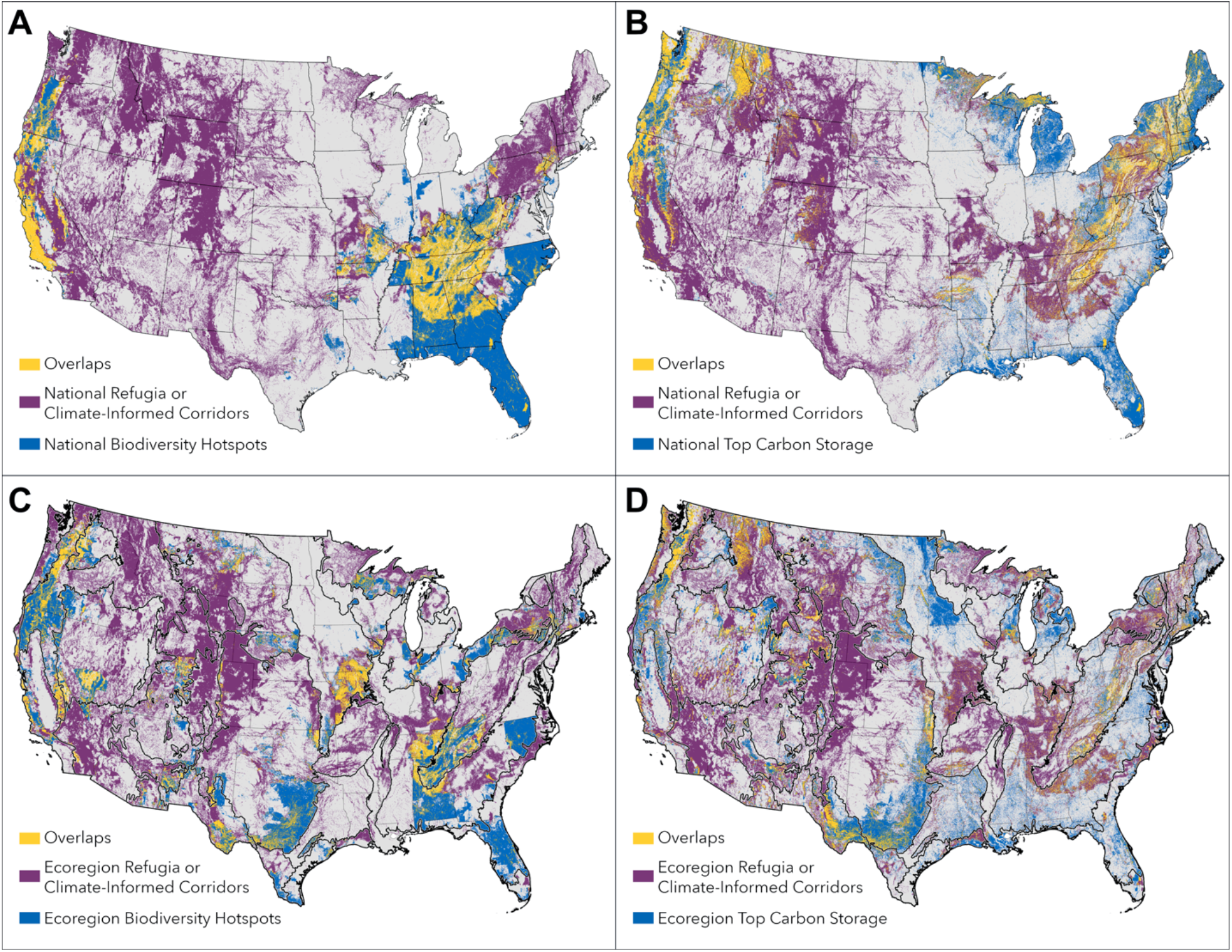
Overlap between national-scale (A,B) and ecoregion-scale (C,D) refugia and corridors with carbon stocks (B,D) and biodiversity hotspots (A,C). The full raster datasets were used to identify refugia in national analyses. Ecoregion-specific analyses employ a stratified approach, where refugia are identified for each ecoregion separately before combining them together. Ecoregions outlined in black in maps C and D.

Taking an ecoregion-specific approach to comparing refugia, corridors, biodiversity, and carbon results in less coincidence: 22.0% and 21.7% of stratified refugia overlap with ecoregion-specific biodiversity hotspots and carbon-rich areas, and 17.5% and 26.1% of corridors overlap with ecoregion-specific biodiversity hotspots and carbon-rich areas, respectively (Fig. 3c& d; Table 3).

**Table 3.**
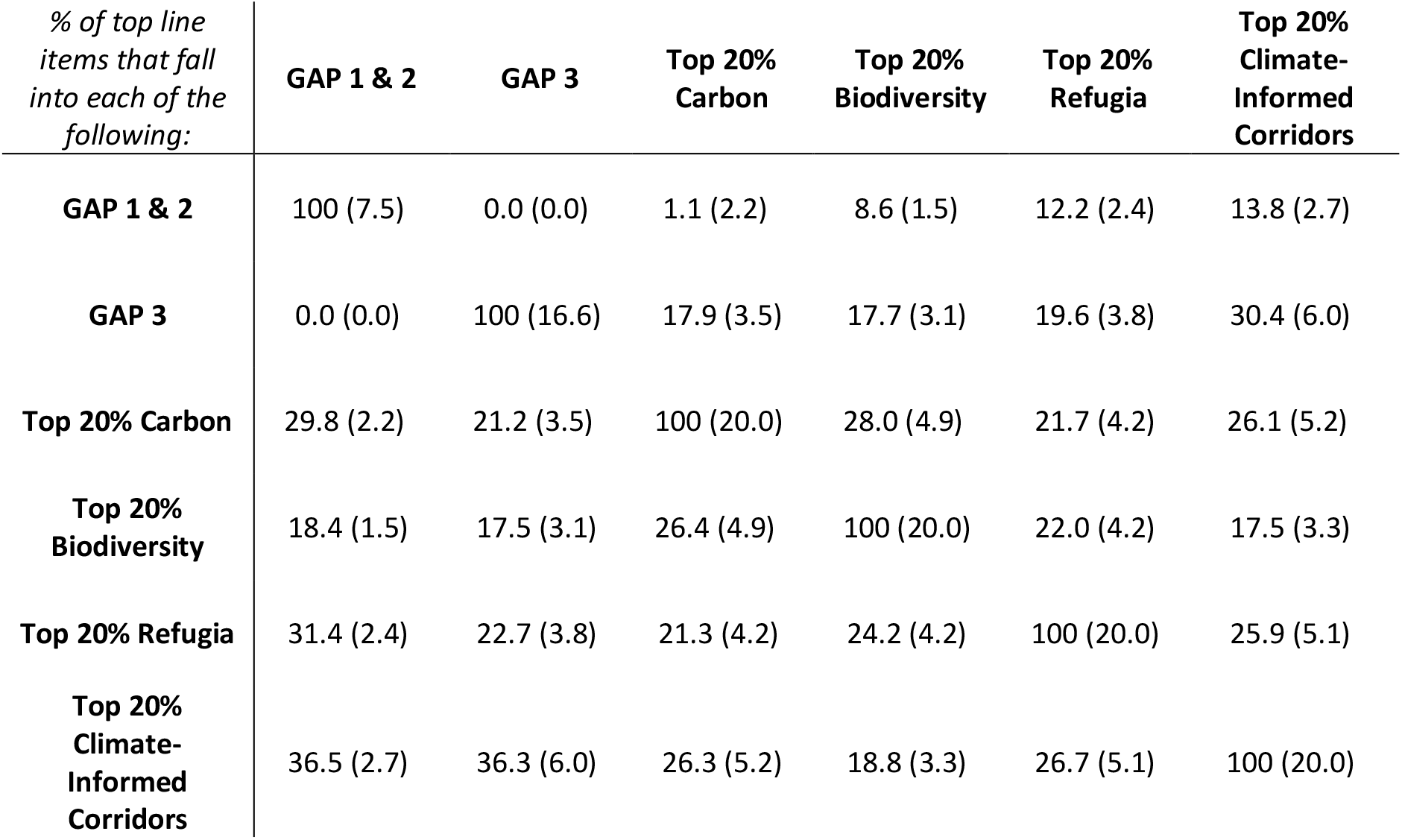
Overlays of ecoregion-specific datasets representing protected areas, carbon stores, biodiversity, climate refugia, and climate corridors. Values represent the percent of each top line item (column) that falls within each row. Values in parentheses are the percent of total CONUS area represented by the overlay.

### Current protections for refugia and corridors

Overall, 12.5% of the combined network of refugia and corridors is managed consistently with biodiversity conservation (i.e., GAP 1 or 2; 4.2% of CONUS or approx. 325,000 km^2^; Fig. 4). The rest of this network falls on GAP 3 (26.5%) or GAP 4 (69.3%) lands, which represents 29.2% of CONUS (approx. 2,280,000 km^2^). Proportions are similar when analyzing protection of national-scale climate refugia and corridors separately (Table 2). Ecoregion-specific refugia fall more heavily in GAP 4 categories with 12.2% of area on lands managed for biodiversity conservation and 19.6% on those managed for multiple uses (Fig. 4, Table 3). Finally, the entire set of CONUS lands representing either biodiversity conservation (GAP 1 or 2) or 30×30 objectives (biodiversity hotspots and/or carbon-rich areas) coincides with 44.5% of the national climate refugia and corridor network.

**Figure 4.**
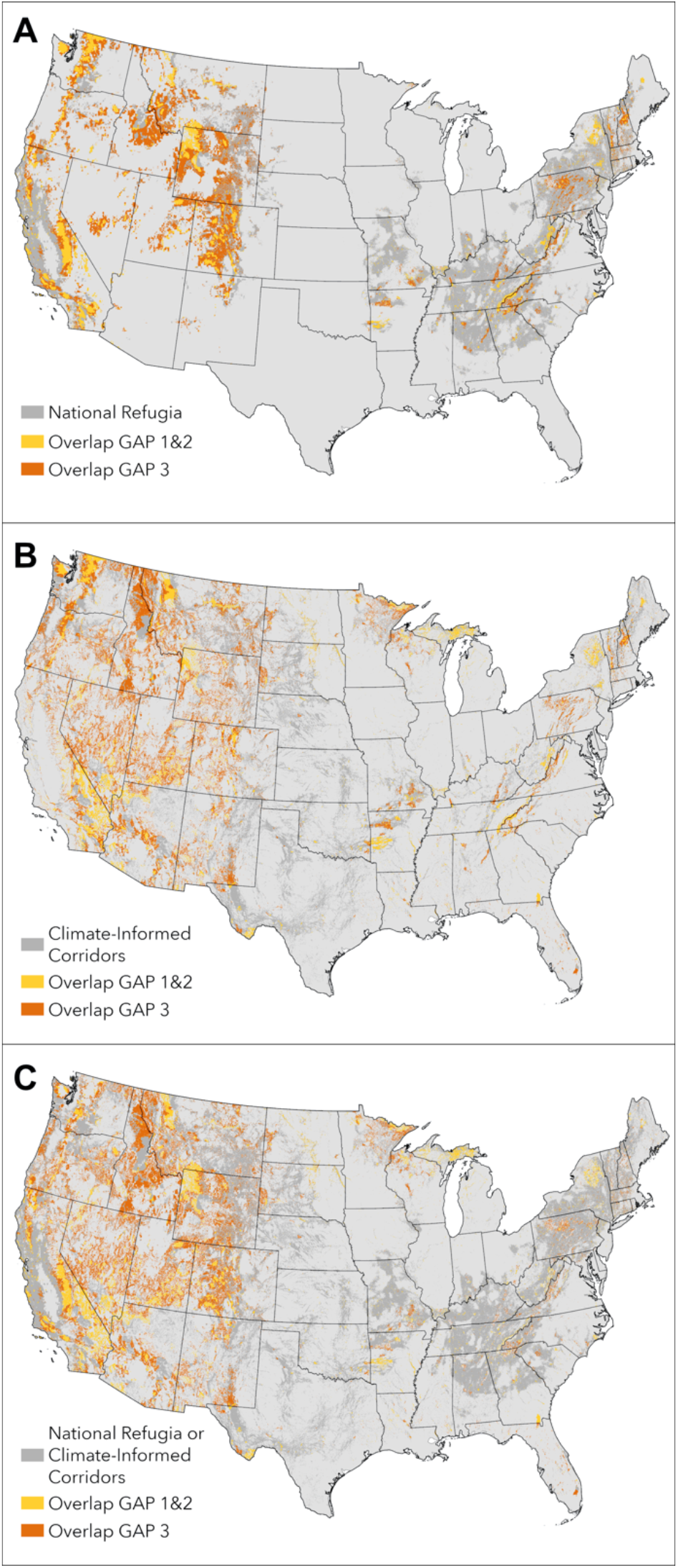
Overlap between national-scale refugia (A), climate corridors (B), and either refugia or corridors (C) with the protected areas database of the US (PADUS). GAP codes are specific to the management intent to conserve biodiversity; GAP 1 and 2 areas are managed in ways typically consistent with conservation and GAP 3 areas are governed under multiple-use mandates that may include biodiversity priorities but may also include incompatible activities.

## DISCUSSION

Currently, the U.S. protected areas network and emerging conservation policy objectives largely fail to represent valuable climate refugia and corridors. While there is some overlap, solely using recent imperiled species ranges and carbon stores as conservation criteria will not inherently protect climate-resilient lands. In the most protective situation - if all biodiversity hotspots and carbon-rich areas were to be considered for strong conservation mandates (e.g., GAP 1 or 2 protections) - a majority (55.5%) of identified climate refugia or corridors would still be left unprotected. The omission of landscapes for climate adaptation from planning initiatives could inhibit the potential for longer-term conservation successes. While simply protecting currently biodiverse or carbon-rich areas may not ensure the preservation of climate corridors and refugia, conserving corridors and refugia will benefit imperiled species in biodiversity-rich hotspots and promote carbon sequestration. This is particularly true in parts of the country (e.g., Appalachia and western California) where hotspots are not directly covered by climate corridors, but adjacent to them, providing opportunities for migration to refugia or future climate analogs.

With over half of the contiguous U.S. identified as at least one type of climate refugia (topodiversity, climatic stability, or tree macrorefugia), many opportunities exist for decision makers interested in future-minded conservation. Like previous work, we demonstrate trade-offs in using one refugia dataset over others: topodiversity data favor environmentally complex regions like mountain ranges, whereas climatic exposure and tree macrorefugia highlight other lands (Michalak et al. 2020). Through our ensemble approach to refugia identification we both highlight the complementary information provided by these approaches (Belote et al. 2018) and simplify varied complex datasets for greater interpretability. A weighted combination of the datasets puts less pressure on the user to choose between mechanisms and on the decision maker to have a deep understanding of the methodology when interpreting maps. However, clarification of a specific refugia type may help states or local municipalities working to set priorities for contributing to national refugia protections based on local environments and community needs.

In addition, taking a combined approach results in high overlap with an ecoregion-stratified approach, suggesting representation of nearly all ecoregions in national efforts focused on conserving climate refugia.

Currently unprotected climate refugia and corridors represent 29.2% of CONUS, of which 38% is federally managed. Given the extent and distribution of land managers, protecting valuable climate adaptation areas can help contribute to the 30% target numerically and meaningfully. However, there will need to be a concerted effort by land managers in all jurisdictions and leadership across jurisdictional boundaries.

### Lands Administered by Government and Tribal Entities

Public lands can make significant contributions to achieving 30×30. The federal lands estate is particularly expansive (20% of CONUS, 86% of PADUS; CRS 2020, Rosa and Malcom 2020) and federal land management agencies are required to varying degrees to prioritize wildlife and habitat conservation. Currently, the majority (86%, representing 18.4% of CONUS) of GAP 3 lands are managed by federal agencies, suggesting that substantial gains can be made in focusing on existing statutory authorities to advance climate-smart conservation on these lands. Of GAP 3 lands, over half are managed by the Bureau of Land Management (BLM) and another third by the U.S. Forest Service (Rosa and Malcom 2020). Both agencies are guided by multiple use management mandates that empower them to designate and manage lands to enhance protection of areas recognized as having important conservation values (respectively, the Federal Land Policy and Management Act of 1976, National Forest Management Act of 1976). The agencies can capitalize on existing land and water designation authorities - like wilderness designation and BLM “areas of critical environmental concern” - to increase protection for climate refugia and corridors.

Expansion of GAP 1 and 2 lands to cover more refugia and corridors can also ensure greater conservation for climate adaptation. The U.S. Fish and Wildlife Service manages the National Wildlife Refuge System (NWRS) expressly to conserve species and habitat, providing a high level of federal land protection. Pursuing the acquisition of lands fundamental to species’ survival and sustainability, including climate refugia and climate corridors, to establish new refuges would be consistent with the purview of NWRS. However, since federal land acquisition and management decisions are often politically contentious, this may be a less feasible option for conserving the additional 440 million acres of land needed to reach the 30% target.

State governments also manage significant acreage (approximately 4% of the U.S.), including state forests, wildlife management areas, game lands, and natural area preserves. State parks, or portions thereof, may also contribute to conservation refugia and corridors, but are often categorized as GAP 4 (i.e., absent or unknown mandates for conservation). States can contribute to 30×30 by upgrading GAP status and management of undeveloped state lands that can further climate adaptation. The State Wildlife Action Planning (SWAP) process also requires each state is to describe “locations and relative condition of key habitats and community types essential to conservation of species” (USFWS & AFWA 2017). Results from this and other studies can help inform this process and help states increasingly update their SWAPs to include climate changes (NFWPCAN 2021).

Tribal nations hold over 56 million acres in trust by the Bureau of Indian Affairs and may manage their lands in ways that afford more substantive protections for lands and species given their lower rates of habitat modification (Lee-Ashley et al. 2019). A long history of managing and observing their lands has provided many indigenous communities with valuable knowledge and experience to inform land management and planning for climate adaptation and resilience (BIA 2018). Respectful inclusion of indigenous systems of knowledges and perspectives “can inform our understanding of how the climate is changing and strategies to adapt to climate change impacts” (NFWPCAN 2021). As such, government-to-government relationships will be important in addressing climate adaptation needs for species and peoples and may include cross-landscape management, tribal involvement in federal and state planning, and more. The Landscape Conservation Cooperative (LCC) program developed by Interior offers one such mechanism to advance landscape-scale protections and coordinate climate-related land conservation activities among Tribal Nations, federal agencies, state, local, and tribal governments, and other stakeholders (NASEM 2016).

### Private and Non-Governmental Organization Lands

As most land in the U.S. is privately owned, conservation efforts on private lands will be critical to expanding protected areas. 62% of the refugia and 56% of corridors fall outside of the protected areas network (GAP 4), but this only represents 20% of CONUS. This suggests that well-targeted, voluntary acquisitions and easements could translate to large gains in private lands conservation. Land trusts are uniquely positioned to scale-up conservation on private lands to achieve the 30×30 target.

In addition to the role of land trusts, private working lands and associated conservation programs can be important to achieving 30×30 (Garibaldi et al. 2020, American Farmland Trust 2021). For instance, Farm Bill programs administered by the U.S. Department of Agriculture such as the Agriculture Conservation Easement Program (ACEP) could be targeted to lands identified as climate refugia or connectivity areas and specify sensitive wetland habitats and riparian areas as eligible lands for wetland easements, as these will be increasingly valuable for supporting wildlife and ecosystem services as the climate changes (Theoharides 2014, Lewis et al. 2019). Longer-term (30 year) contracts that offer a commitment to re-enrollment should be encouraged to ensure enduring conservation measures. Additionally, Environmental Quality Incentives Program (EQIP) and the Conservation Stewardship Program (CSP) can better reflect climate adaptation needs by assigning higher ranking to practices designed to build resilient landscapes (Theoharides 2014).

### Limitations

To enhance species’ resilience in the face of growing climate and biodiversity crises, corridors and refugia must be preserved across both lands and waters. Due to some limitations of data and our analyses, we recommend against siting protections based on the coincidence of current biodiversity/carbon hotspots and climate refugia/corridors alone. For one, complementarity of species assemblages is not accounted for and there may be biases toward conserving certain taxa. Additionally, while we include aquatic species in our biodiversity metric, and wetland/riparian areas in some topographic measures of refugia, we did not explicitly include aquatic refugia. Currently, there is no complete national dataset to represent aquatic refugia. Because cold-water aquatic organisms are among the most vulnerable taxa to climate change, future analyses should focus on identifying freshwater refugia and corridors where sufficient data exists. Given the international scope of 30×30 and the benefits of larger-scale connectivity, future work on climate adaptation in 30×30 implementation should look beyond terrestrial habitats and political boundaries to cover all ecosystems of North America.

Our analysis demonstrates the need to make climate adaptation a more explicit objective in conservation planning for addressing the biodiversity crisis. Without direct consideration for climate refugia and corridors, a 30×30 implementation focused on current species ranges and carbon stocks may be ineffective for the longer-term persistence of species. The key to operationalizing 30×30 and subsequent efforts will be growing a protected areas network that ensures a long-term commitment to biodiversity and climate. By incorporating climate refugia and corridors, the U.S. can work to protect places that will continue to serve wildlife and human populations now and in the future.

## ACKNOWLEDGEMENTS

We thank M. Anderson, J. Grand, J. Lawler, R. List, J. Michalak, S. Saunders, and R. Wynn-Grant for their thoughtful review of the project and engaging discussion over key concepts. Additional gratitude goes to those organizations that make these datasets publicly available to enable this and other research.

Recent work in the field of conservation science and STEM at large has identified a bias in citation practices such that papers from women and minorities are relatively under-cited (see Rudd et al. 2021, Larivière et al. 2013, and others). While we did not proactively choose references that reflect the diversity of the field in thought, form or contribution, gender, and other factors for this work, we recognize the biases that may have been unintentionally introduced. We look forward to future work that can help us to better understand how to support equitable practices in conservation science.

## DATA AVAILABILITY

The data that support the findings of this study are all publicly available. Those directly generated by this work can be downloaded from the following OSF repository: https://osf.io/jksyx/

