## Supplemental Materials for "Targeting current species ranges and carbon stocks fails to conserve biodiversity in a changing climate: opportunities to support climate adaptation under 30×30"

**Table S1.**  Correlation matrix across all eight refugia datasets.

|  | **Bird Macrorefugia** | **Climatic Dissimilarity** | **Current Climate Diversity** | **Ecotypic Diversity** | **Climate Velocity** | **Landscape Diversity** | **Land Facet Diversity** | **Tree Macrorefugia** |
| --- | --- | --- | --- | --- | --- | --- | --- | --- |
| **Bird Macrorefugia** | 1.000 | 0.175 | 0.565 | 0.339 | -0.069 | 0.093 | 0.446 | 0.233 |
| **Climatic Dissimilarity** | 0.175 | 1.000 | 0.159 | 0.166 | 0.243 | 0.048 | 0.162 | 0.019 |
| **Current Climate Diversity** | 0.565 | 0.159 | 1.000 | 0.460 | -0.221 | 0.177 | 0.572 | 0.047 |
| **Ecotypic Diversity** | 0.339 | 0.166 | 0.460 | 1.000 | -0.134 | 0.174 | 0.570 | 0.008 |
| **Climate Velocity** | -0.069 | 0.243 | -0.221 | -0.134 | 1.000 | -0.054 | -0.192 | 0.042 |
| **Landscape Diversity** | 0.093 | 0.047 | 0.177 | 0.174 | -0.054 | 1.000 | 0.253 | -0.043 |
| **Land Facet Diversity** | 0.446 | 0.162 | 0.572 | 0.570 | -0.192 | 0.253 | 1.000 | 0.178 |
| **Tree Macrorefugia** | 0.233 | 0.019 | 0.047 | 0.008 | 0.042 | -0.043 | 0.178 | 1.000 |

**Table S2.**  Eigenvectors for all eight refugia datasets across the first three principal components. Grays indicate dataset groupings that define refugia classes (see Table 1). Datasets aligned with component one represent topodiversity, component two represents climate stability, and component three represents tree macrorefugia. Generally, datasets were aligned with a component based on the sign and size of their eigenvector. For the landscape diversity dataset, sign rather than strength of the eigenvector was chosen to avoid tradeoff with other datasets (namely, tree macrorefugia) within the refugia class. Together these components explain 63.4% of the variance.

|  | **Principal Component 1** | **Principal Component 2** | **Principal Component 3** |
| --- | --- | --- | --- |
| **Bird Macrorefugia** | **0.443** | 0.160 | 0.242 |
| **Climatic Dissimilarity** | 0.168 | **0.618** | -0.348 |
| **Current Climate Diversity** | **0.496** | -0.075 | -0.017 |
| **Ecotypic Diversity** | **0.441** | -0.051 | -0.196 |
| **Climate Velocity** | -0.153 | **0.691** | -0.167 |
| **Landscape Diversity** | **0.193** | -0.128 | -0.405 |
| **Land Facet Diversity** | **0.506** | -0.042 | 0.006 |
| **Tree Macrorefugia** | 0.126 | 0.298 | **0.768** |

**Table S3.**  Covariance matrix across all eight refugia datasets.

|  | **Bird Macrorefugia** | **Climatic Dissimilarity** | **Current Climate Diversity** | **Ecotypic Diversity** | **Climate Velocity** | **Landscape Diversity** | **Land Facet Diversity** | **Tree Macrorefugia** |
| --- | --- | --- | --- | --- | --- | --- | --- | --- |
| **Bird Macrorefugia** | 0.190 | 0.033 | 0.106 | 0.064 | -0.013 | 0.017 | 0.084 | 0.044 |
| **Climatic Dissimilarity** | 0.033 | 0.187 | 0.030 | 0.031 | 0.045 | 0.009 | 0.030 | 0.004 |
| **Current Climate Diversity** | 0.106 | 0.030 | 0.187 | 0.086 | -0.041 | 0.033 | 0.107 | 0.009 |
| **Ecotypic Diversity** | 0.064 | 0.031 | 0.086 | 0.187 | -0.025 | 0.032 | 0.107 | 0.001 |
| **Climate Velocity** | -0.013 | 0.045 | -0.041 | -0.025 | 0.186 | -0.010 | -0.036 | 0.008 |
| **Landscape Diversity** | 0.017 | 0.009 | 0.033 | 0.032 | -0.010 | 0.182 | 0.047 | -0.008 |
| **Land Facet Diversity** | 0.084 | 0.030 | 0.107 | 0.107 | -0.036 | 0.047 | 0.187 | 0.034 |
| **Tree Macrorefugia** | 0.044 | 0.004 | 0.009 | 0.001 | 0.008 | -0.008 | 0.034 | 0.192 |

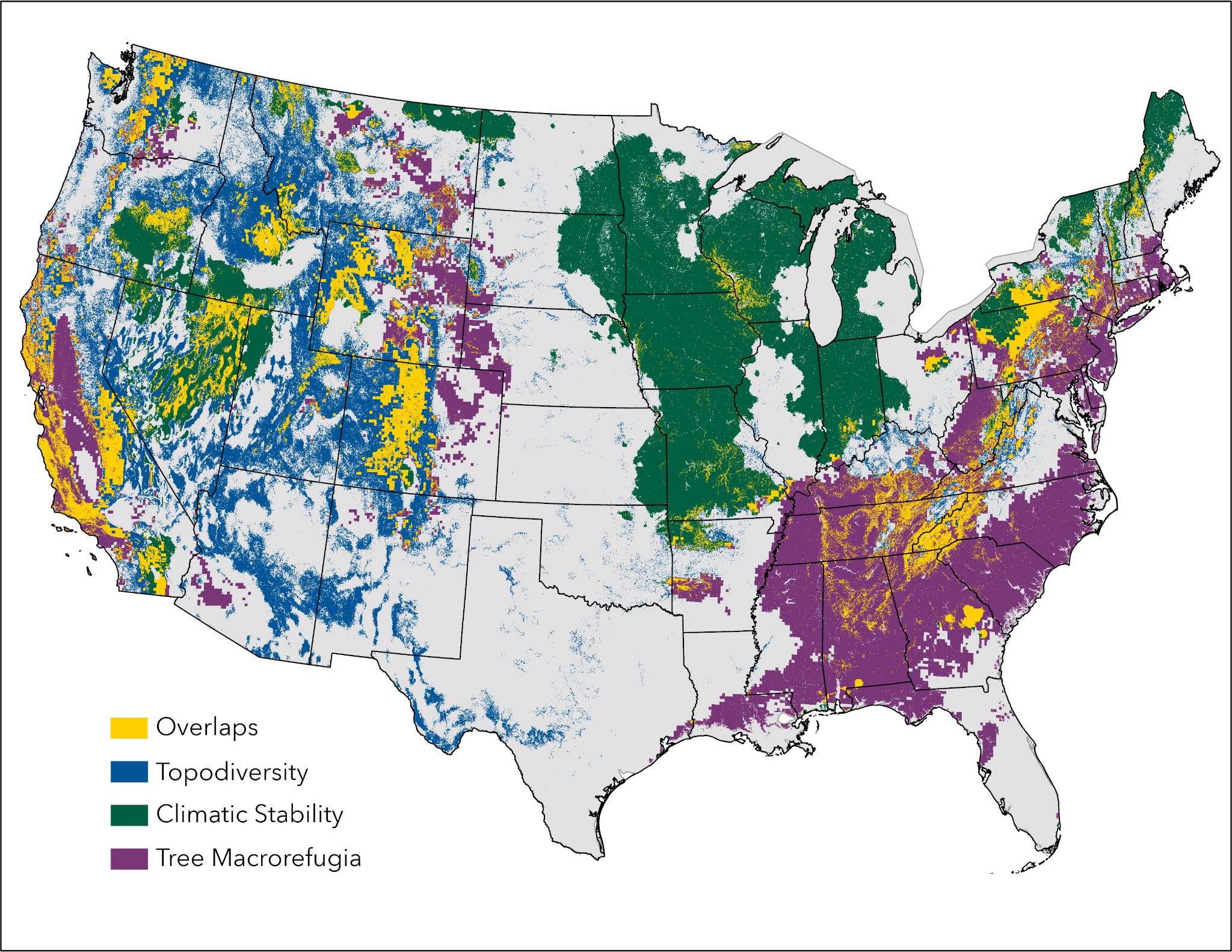

**Figure S1.** Locations falling within the top 20% of the distribution of values for each refugia class. Classes are groupings of refugia datasets and are chosen based on results from a principal components analysis where component 1 (topodiversity, n = 5) explained 33.8%, component 2 (climate stability, n = 2) explained 15.9% and component 3 (tree macrorefugia, n = 1) explained near 13.8% of variation. See Table 1 for dataset descriptions.
